# Inoculation of biocontrol bacteria alleviated *Panax ginseng* replanting problem

**DOI:** 10.1101/143412

**Authors:** Linlin Dong, Jiang Xu, Lianjuan Zhang, Guangfei Wei, He Su, Juan Yang, Jun Qian, Ran Xu, Baosheng Liao, Liang Shen, Mingli Wu, Ruiyang Cheng, Shilin Chen

**Affiliations:** Key Laboratory of Beijing for Identification and Safety Evaluation of Chinese Medicine, Institute of Chinese Materia Medica, China Academy of Chinese Medical Sciences, Beijing 100700, China.

**Keywords:** *Panax ginseng*, microbial communities, plant age, developmental stage, replanting problem, biocontrol

## Abstract

Replanting problem is a common and serious issue hindering the continuous cultivation of *Panax* plants. Changes in soil microbial community driven by plant species of different ages and developmental stages are speculated to cause this problem. Inoculation of microbial antagonists is proposed to alleviate replanting issues efficiently.

High-throughput sequencing revealed that bacterial diversity evidently decreased, and fungal diversity markedly increased in soils of adult ginseng plants in the root growth stage. Relatively few beneficial microbe agents, such as *Luteolibacter*, Cytophagaceae, *Luteibacter, Sphingomonas*, Sphingomonadaceae, and Zygomycota, were observed. On the contrary, the relative abundance of harmful microorganism agents, namely, Brevundimonas, Enterobacteriaceae, *Pandoraea*, Cantharellales, *Dendryphion, Fusarium*, and Chytridiomycota, increased with pant age. Furthermore, *Bacillus subtilis* 50-1 was isolated and served as microbial antagonists against pathogenic *Fusarium oxysporum* of ginseng root-rot, and its biocontrol efficacy was 67.8% using a dual culture assay. The ginseng death rate and relative abundance of *Fusarium* decreased by 63.3% and 46.1%, respectively, after inoculation with 50-1 in replanting soils. Data revealed that changes in the diversity and composition of rhizospheric microbial communities driven by ginseng of different ages and developmental stages could cause microecological degradation. Biocontrol using microbial antagonists was an effective method for alleviating the replanting problem.

**Highlight:** Changes in rhizospheric microbial communities driven by ginseng plants 13 of different ages and developmental stages could cause microecological degradation. 14 Biocontrol using microbial antagonists effectively alleviated the replanting problem.

## Introduction

*Panax ginseng* C.A. Meyer demonstrates anti-inflammatory and antitumor effects, and it is commonly used in traditional Chinese medicine (Choo *et al*., 2008; Ernst, 2010). The current annual global market value of this species is approximately 3.5 billion dollars (Hong *et al*., 2006). Wild ginseng resources are becoming scarce because of excessive and predatory exploitation, and wild ginseng is gradually substituted with cultivated ginseng in mainstream market (Li *et al*., 2012; Wu *et al*., 2013). Ginseng is continuously cultivated in fixed plots for 4–5 years, and subsequent replanting usually fails because of continuous cropping obstacles (Ying *et al*., 2012a). Several years of crop rotation is needed in successful replanting. Large-scale deforestation increases to satisfy the market demand for ginseng. Moreover, conflict between the operation of ginseng industry and environmental conservation has worsened. Replanting issue is a severe drawback hindering the development of ginseng industry. Hence, this problem must be addressed urgently.

Factors such as deterioration of soil physicochemical properties, allelopathy/autotoxicity, outburst of soil-borne disease, and soil microbial community changes cause replanting problem (Ogweno & Yu, 2006; Wu *et al*., 2008; Huang *et al*., 2013). Changes in soil microbial community is one of the main factors hindering crop replantation (Nayyar *et al*., 2009; Manici *et al*., 2013). Xiao *et al*. (2016) reported that the lack of balance in rhizospheric microbial communities is one of the key factors causing discontinuous ginseng cultivation. Microbial communities exhibit evident changes during ginseng cultivation, thereby causing an imbalance in microbial community (Ying *et al*., 2012b). Increase in the relative abundance of pathogenic microorganism could lead to the occurrence of soil-borne disease. Taken together, changes in rhizospheric microbial community are suggested to cause the replanting issues.

Many biotic or abiotic factors lead to changes in rhizospheric microbial communities. Plants of different ages can alter microbial community (Wu *et al*., 2015). Differences in diversity and composition of soil microbial community were observed under continuous cropping of *P. notoginseng;* moreover, fungal diversity and microbial taxa in soil microbial community are related to seedling mortality (Dong *et al*., 2016). Li *et al*. (2012) found that differences between rhizospheric and non-rhizospheric soil microbial community in a particular site tend to become increasingly significant with ginseng growth. *Panax* plants with different ages in a site drive changes in microbial community. Plant developmental stage influences microbial community structure and activity, and marked shifts in diversity and relative activity are observed in soil microbial communities of different developmental stages (Houlden *et al*., 2008). However, the information whether *Panax* plants of different ages and developmental stages mediate microbial community is unclear. We hypothesized that changes in the rhizospheric microbial community driven by ginseng plants of different ages and developmental stages could cause microecological degradation.

Root rot is a severe disease that hinders the replantation of *Panax* plants (Guo *et al*., 2009). This soil-borne disease is caused by the pathogenic fungus *Fusarium oxysporum*, which is the main agent in *Panax* plants (Miao *et al*., 2006; Dong *et al*., 2016).The relative abundance of *F. oxysporum* increased with the notoginseng cultivation and death rate of seedlings was significantly related to its abundance (Dong *et al*., 2016). Thus decrease in the abundance of pathogenic *F. oxysporum* could alleviate the occurrence of root-rot. Biological control using microbial antagonists has attracted interest as an effective method to control plant pathogens due to its non-toxic nature (Zheng *et al*., 2011). Many studies reported that biocontrol bacteria, such as *Pseudomonas fluorescens* and *Bacillus amyloliquefaciens* RWL-1, show evident growth inhibition activity against *F. oxysporum* in tomato (Kamilova *et al*., 2006; Shahzad *et al*., 2017). Nevertheless, microbial antagonists against ginseng root rot is scarce, and application of biocontrol bacteria could be effective to alleviate the replanting problem.

Recently, herbgenomics has been raised for medicinal plants research using genomic tools including investigating rhizospheric environment using metagenomic sequencing (Chen *et al*, 2015). In the present study, high-throughput sequencing analysis of 16S and 18S rRNA genes was used to demonstrate changes in the diversity and composition of soil microbial communities in the rhizosphere of ginseng seedlings of different ages and developmental stages. Furthermore, biocontrol bacteria against *F. oxysporum* were isolated using a dual culture technique, and their inhibition activity in a replanting soil was confirmed. These results increased our understanding of the reasons behind replanting issues caused by rhizospheric microbial community. Data provided an effective method for soil bioremediation to alleviate issues in replanting of Chinese herbal medicine.

## Materials and Methods

### Field experiment and soil extraction

This field experiment was performed in a ginseng plantation in Jingyu, Jilin Province (42°20☐N, 126°50☐E, 775 m a.s.l.), which is the main ginseng-producing region in China. This region receives a northern temperate continental climate and an annual precipitation of approximately 767 mm. The plough layer in the plantation consists of gray brown soil.

Ginseng seedlings are consecutively grown for 4–5 years in a fixed site before their root can be harvested. The disease and mortality of ginseng seedlings generally occur after being consecutively grown for 2 years. Thus, we analyzed the influence of two-, three-, and four-year-old transplanted seedlings on rhizospheric microbial communities. In this experiment, two-, three-, and four-year-old ginseng seedlings were transplanted in each plot in our plantation and denoted as 2-y, 3-y, and 4-y, respectively. In our plantation, field plots were arranged in a completely randomized block design, and three replicate plots (1.7 m × 8.0 m) were used for each plant age. Ginseng was cultivated strictly in accordance with the standard operating procedures of good agricultural practice (Heuberger *et al*., 2010; Zhang *et al*., 2010).

The distinct stages of ginseng development are as follows: vegetative, flowering, fruiting, root growth, and annual dormancy (Table S1). During dormancy, the aboveground plant parts of ginseng wither, and the root becomes dormant. In this stage, the underground root activities weaken, and soil samples obtained during this stage are not included in analyses. This experiment included 36 soil samples obtained from two-, three-, and four-year-old ginseng seedlings under four developmental stages, namely, vegetative (2-Ve, 3-Ve, and 4-Ve), flowering (2-Fl,3-Fl, and 4-Fl), fruiting (2-Fr, 3-Fr, and 4-Fr), and root growth (2-Ro, 3-Ro, and 4-Ro). Six ginseng seedlings were randomly obtained from each plot (1.7 m × 8.0 m); the soils were removed, and rhizosphere fractions were brushed and pooled into one sample. Soil samples were obtained from three replicates per treatment. Thirty-six soil samples were homogenized by passing through a 2 mm sieve for further processing. The characteristics of the soil samples are described in Table S2.

### Calculation of emergence and survival rates

The ginseng emergence rate was analyzed in May. The emergence rate in each plot was calculated as follows: number of emerging ginseng seedlings divided by the total number of transplanted seedlings. The mortality of ginseng seedlings was observed from June to August, and the survival rate was determined at the end of August. The survival rate of ginseng in each plot was calculated as follows: number of living seedlings divided by the total number of emerging seedlings in the area. Three plots per treatment served as three replicates.

### DNA extraction and PCR amplification

Total soil DNA was extracted from 0.1 g of freeze-dried soil using a MoBio Powersoil Kit (MoBio Laboratories Inc., Carlsbad, CA) according to the manufacturer’s instructions; the DNA was stored at –20 °C for further processing. For each sample, fragments of 16S and 18S rRNA genes were amplified using the conserved primers 27F/338R (Fierer *et al*., 2008) and 817F/1196R (Rousk *et al*., 2010). The forward and reverse primers contained an eight-base pair barcode (Table S3). Amplification reactions and purification were performed as previously described (Rodrigues *et al*., 2013). Purified PCR products were quantified by Qubit®3.0 (Life Invitrogen, Germany), and these amplicons were pooled in equimolar ratios for sequencing.

### High-throughput sequencing

The pooled DNA product was used to construct Illumina pair-end library according to the Illumina’s genomic DNA library preparation procedure. The amplicon library was subsequently paired-end sequenced (2 × 250) on an Illumina HiSeq platform (Shanghai Biozeron Co., Ltd.) according to standard protocols.

Raw FASTQ files were demultiplexed and quality-filtered using QIIME with the following pipeline (Caporaso *et al*., 2010). Operational taxonomic units were clustered using 97% similarity cutoff by using UPARSE (version 7.1 http://drive5.com/uparse/), and chimeric sequences were identified and removed using UCHIME. The phylogenetic affiliation of each 16S and 18S rRNA gene sequence was analyzed using Ribosomal Database Project (Wang *et al*., 2007) and Silva schemes (Quast *et al*., 2013). Rarefaction analysis based on Mothur v. 1.21.1 (Schloss *et al*., 2009) was performed to reveal diversity indexes, including Chao 1 and Shannon diversity (*H*☐) indexes. We clustered the taxa obtained from the RDP Classifier through complete linkage hierarchical clustering by using *R* package HCLUST (http://sekhon.berkeley.edu/stats/html/hclust.html). PCoA was used to compare groups of samples based on unweighted UniFrac distance metrics in QIIME (Caporaso *et al*., 2010). Linear discriminant analysis effect size (LEfSe) (http://huttenhower.sph.harvard.edu/lefse/) was used to characterize the features differentiating the bacterial and fungal communities in soils according to the methods described by Segata *et al*. (2011). All metagenomic data were submitted to the National Center for Biotechnology Information (http://www.ncbi.nlm.nih.gov); the accession numbers of 16S rRNA and 18S rRNA genes are SRP103168 and SRP103176, respectively.

### Quantitative PCR of *Fusarium*

To compare the relative abundance of *Fusarium* in soils of ginseng seedlings, the copy numbers of *Fusarium* were calculated using the ITS-Fu-F/ITS-Fu-R (Abd-Elsalam *et al*., 2003). Quantitative PCR was performed according to the description of Rousk *et al*. (2010) with minor modification. Standard curves were generated using 10-fold serial dilution of a plasmid containing a full-length copy of *F. oxysporum* 18S rRNA gene to estimate the copy numbers of *Fusarium*. qPCR reactions (25 μL) were performed using a SYBR Green PCR Master mix (Takara, Toyobo, Japan). *Fusarium* copy numbers were generated using a regression equation for each assay relating the cycle threshold value to the known number of copies in the standards.

### Isolation and selection of antagonistic bacteria

Biological control is one of the most remarkable potential approaches for plant disease control. Due to the increase in the abundance of pathogenic agents of ginseng root rot, a dual culture assay was used to screen microbial antagonists against *F. oxysporum*. Rhizosphere soils of three-year ginseng seedlings under root growth stage were used to isolate and select the antagonistic bacteria against *F. oxysporum*. The pathogenic *F. oxysporum* was isolated and confirmed in our previous study (Wang *et al*., 2016). Soil (10 g) was homogenized in 100 mL of sterile distilled water, and the bacteria were isolated through serial dilution technique. The isolated single strain was screened on the basis of its antagonistic activity against *F. oxysporum* in a dual culture according to the description of Shahzad *et al*. (2017). The experiment was replicated three times. The zone of inhibition was measured following the description of Kaiser *et al*. (2005) to screen the antagonistic bacterium and examine the antagonistic activity of the candidates.

### Identification of antagonistic bacterial strain

Antagonistic bacterium (named strain 50-1) was identified by morphological and molecular methods. The morphology of strain 50-1 was recorded after it was incubated on Luria–Bertani (yeast extract (5 g), peptone (10 g), NaCl (10 g), and agar (10 g) in 1 L of water) medium for 24 h later. Strain 50-1 was molecularly identified by amplifying 16S rRNA according to the description by Cai *et al*. (2012). The amplified PCR product was analyzed on a 3730 XL sequencer (Applied Biosystems, Foster City, CA, USA), and the generated sequence (accession number KY962803) was submitted to GenBank. Neighbor-joining trees were constructed in MEGA v6.0 to generate Kimura 2-parameter distance matrixes for each sequence following standard parameters. The numbers at the branch knots were bootstrap values based on 1000 resamplings for the maximum likelihood.

To further identify the strain 50-1, total genomic DNA from strain 50-1 was extracted (Tiangen, Beijing, China) and purified by RNase-free DNase I (Takara, Kyoto, Japan) to analyze its genome. The complete genome sequence was assembled using a hybrid sequencing strategy combining PacBio RS II and Illumina HiSeq sequencing platforms (Koren *et al*., 2012). Genome sequencing was performed with Illumina HiSeq2500 using the PE250 strategy following the manufacturer’s protocol. The reads obtained with Illumina PCR adapter and the filtered low-quality reads were assembled by SOAPdenovo (Li *et al*., 2008; Li *et al*., 2010) to generate scaffolds. Gene prediction genome assembly was performed using Glimmer and gene functions were annotated by BLASTP against NR, COG and KEGG databases. GeneMarkS (Besemer *et al*., 2001) with an integrated model combining the GeneMarkS-generated parameters and Heuristic model parameters. A genome overview with annotation information was created using Circos (Krzywinski *et al*., 2009). The whole genome was deposited in NCBI (BioProject ID PRJNA383782).

### Evaluation of the biocontrol efficacy of antagonists in replanting soils

A pot experiment was performed to assess the biocontrol efficacy of bacterium 50-1 in replanting soils in our phytotron. Each pot contained 1 kg of soils cultivated with ginseng seedlings for three years and happened root rot. Two-year-old ginseng seedlings (three plants) were transplanted in each pot. The pots were placed in the phytotron under the following conditions: 26±2 °C, 60% humidity, and 14 h of light alternating with 10 h of darkness. The rhizospheres of uniform ginseng seedlings were inoculated with 1 mL (10^6^ cfu mL–^1^) of cultures containing biocontrol strains when the seedlings were cultivated for one week. Pots were inoculated with cultures containing inactivated strains as controls. Five pots served as one replicate, and three replicates were prepared. After a two months following inoculation, the death rate of ginseng seedlings was calculated as follows: number of dead seedlings divided by the total number of transplanted seedlings in each treatment. The rhizosphere soils of three seedlings were randomly selected from each treatment, and they served as one sample for the analysis of the relative abundance of *Fusarium*. The experiment was replicated three times.

## Statistical analyses

SPSS version 16.0 software (SPSS Inc., Chicago, IL, USA) was used for statistical analyses. Variables were considered for all treatment replicates and subjected to ANOVA. Mean values were compared by calculating the least significant di☐erence at 5% level.

## Results

### Emergence and survival rates of ginseng seedlings

The ginseng plants grown on field showed a healthy appearance with no sign of disease during the growing season (Figs. 1A–1C). The emergence and survival rates exhibited no significant difference among two-, three-, or four-year-old ginseng seedlings (Figs. 1D, 1E). The emergence rate of two-, three-, or four-year-old ginseng seedlings ranged from 82.3% to 86.9% (Fig. 1D), and their survival rate was 94.1%–97.1% (Fig. 1E). The growth of ginseng seedlings of different ages was favorable, which guaranteed the success of the field experiment.

**Figure 1.**
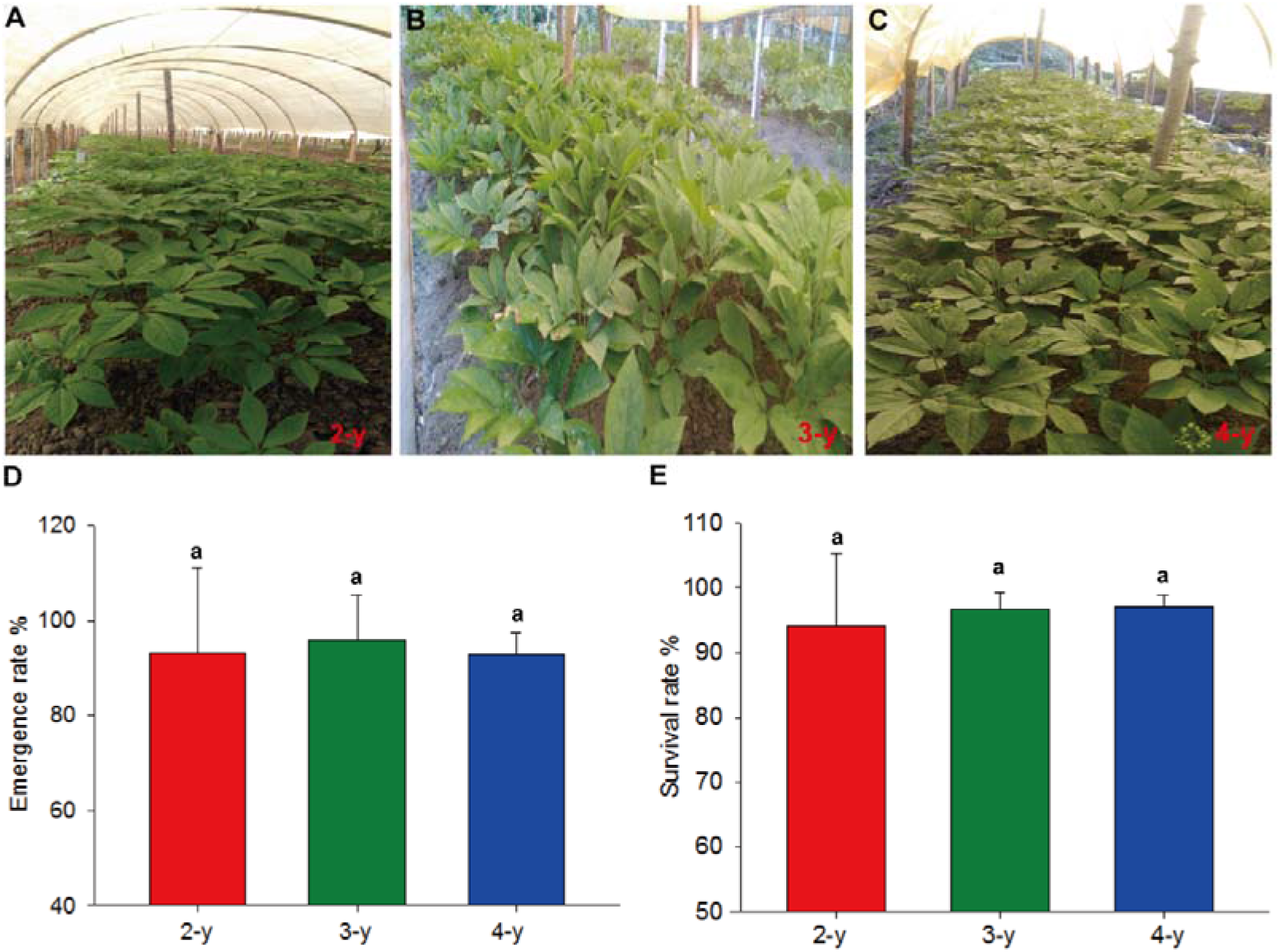
Growth status of ginseng seedlings. (A–C) Two-, three-, and four-year-old ginseng seedlings, respectively. (D) Emergence rate of ginseng seedlings. (E) Survival rate of ginseng seedlings. 2-y, 3-y, and 4-y represents the transplanted two-, three-, and four-year-old seedlings, respectively. Data are presented as mean ± SD (*n* = 3). Identical letters denote nonsignificant difference at 0.05 significance level.

### Taxonomic diversity of bacterial and fungal communities

A total of 2,296,684 classifiable 16S rRNA sequence reads were obtained from 36 soil samples for analysis (Table S3). The mean number of classifiable sequences per sample was 63,796 (dominant length: 283–293 bp). The bacterial diversity indexes (*H*’ and Chao 1) markedly changed in the rhizosphere of ginseng seedlings of different ages in late developmental stages (Fig. 2). *H*’ and Chao 1 values were significantly higher in the rhizosphere of 4-y seedlings than those in the rhizosphere of 2-y seedlings in fruiting stage (Figs. 2A, 2B). During root growth, *H*’ and Chao 1 values evidently decreased in the rhizospheres of 3-y and 4-y seedlings than those in the rhizosphere of 2-y seedlings.

**Figure 2.**
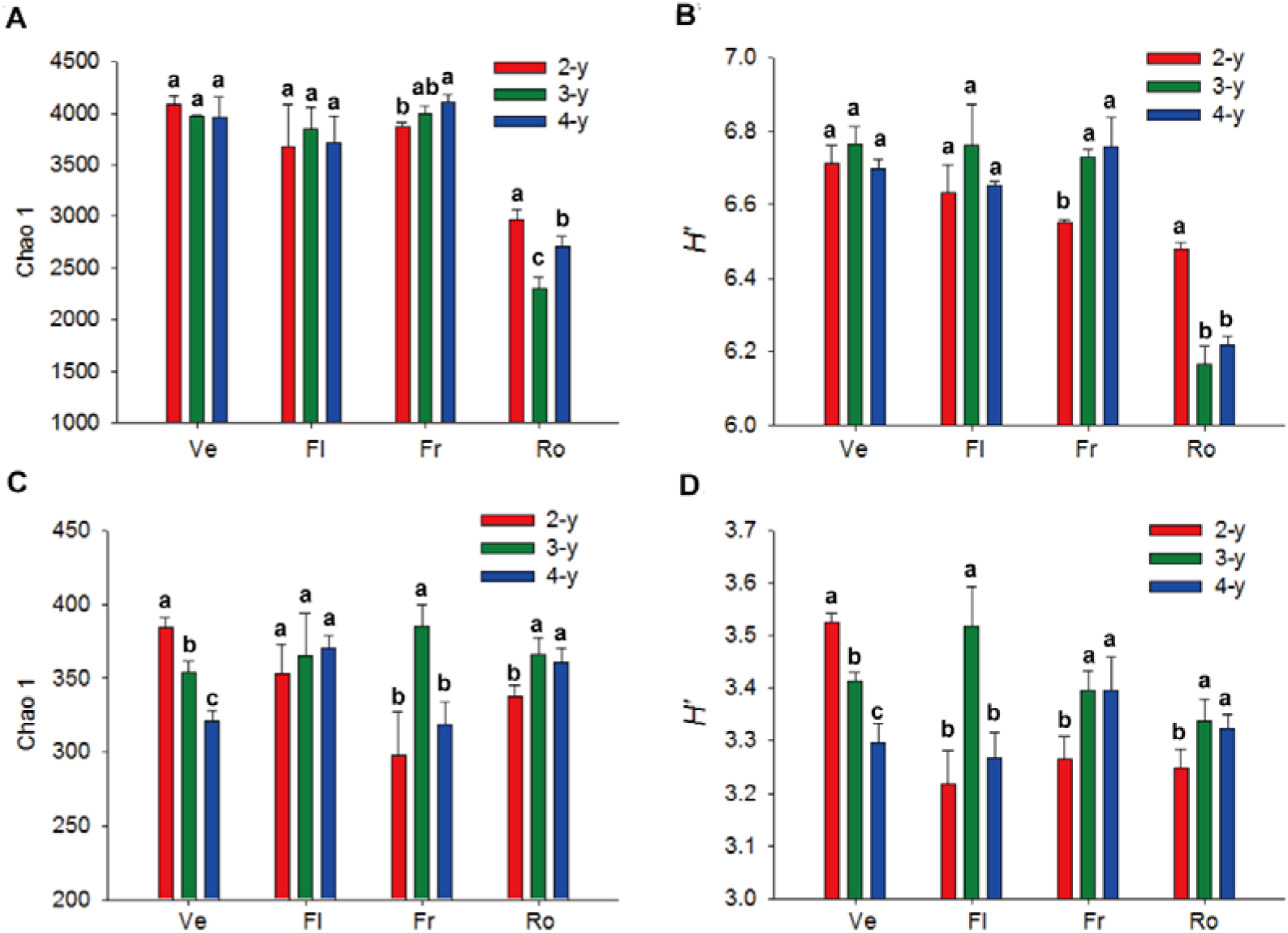
Bacterial and fungal diversities in the rhizosphere of ginseng seedlings of different ages and developmental stages. (A, B) show the Chao 1 and Shannon diversity (*H*) of bacterial community. (C, D) show the Chao 1 and *H*’ of fungal community. Ve, Fl, Fr, and Ro represent vegetative, flowering, fruiting, and root growth stages, respectively. Data are presented as mean ± SD (*n* = 3). Non-identical letters denote significant difference in the same developmental stage of ginseng plant with different ages at 0.05 significance level.

A total of 1,321,521 classifiable fungi sequences with a mean of 36,709 sequences were obtained per soil sample (dominant length: 411–414 bp; Table S3). *H*’ and Chao 1 values markedly decreased in the rhizosphere with increasing ginseng age under vegetative stage (Figs. 2C, 2D). During root growth, *H*’ and Chao 1 values were evidently higher in the soils of 3-y and 4-y seedlings than those in the soil of 2-y seedlings.

### Variation in bacterial and fungal community compositions

PCoA ordination and Bray–Curtis distance matrix revealed evident difference in bacterial communities in soils of ginseng plant of different ages and developmental stages (Figs. 3, S1). In vegetative and flowering stages, the second principal components (14.41% and 14.22% contributions, respectively) demonstrated that the bacterial communities in soils of 2-y seedlings differed from those in soils of 4-y seedlings (Figs. 3A, 3B). In fruiting stage, the first principal component axis (22.00% contributions) indicated that the bacterial communities in soils of 2-y seedlings significantly differed from those in soils of 3-y seedlings (Fig. 3C). Additionally, the first principal component axis (22.29% contributions) indicated that the bacterial communities in soils of 3-year-old seedlings significantly differed from those in soils of 2-y and 4-y seedlings; furthermore, the second principal component (13.98% contributions) demonstrated that the bacterial communities in soils of 2-y seedlings differ from those in soils of 4-y seedlings (Fig. 3D). LEfSe revealed differences in the rhizospheric bacterial communities of ginseng seedlings of different ages and developmental stages (Fig. S2).

**Figure 3.**
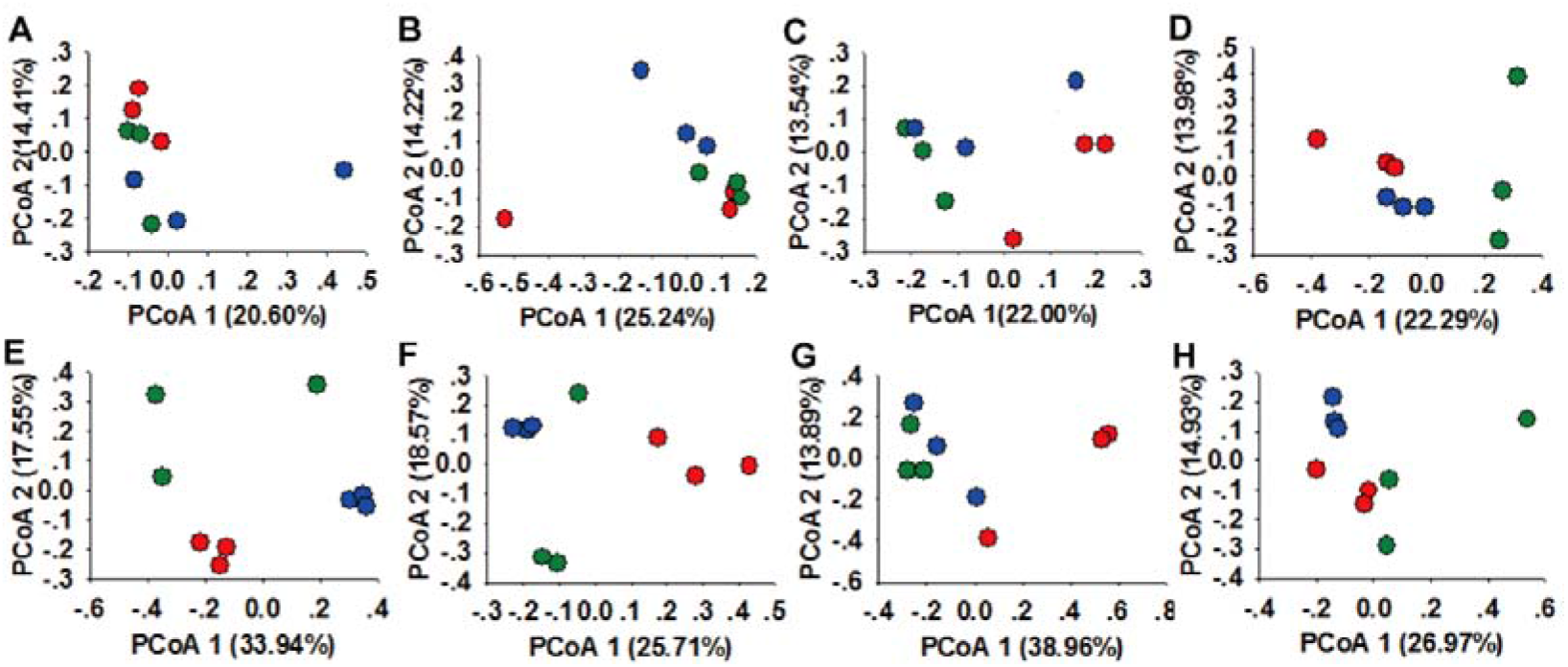
Changes in bacterial and fungal communities in the rhizosphere of ginseng seedlings of different ages and developmental stages. (A–D) PCoA ordination plots display the relatedness of samples separated using Unweighted UniFrac distance of classified 16S rRNA gene sequences at vegetative, flowering, fruiting, and root growth stages, respectively. (E–H) PCoA ordination plots show the relatedness of samples separated using Unweighted UniFrac distance of classified 18S rRNA gene sequences at the Ve, Fl, Fr, and Ro stages, respectively. Red, green, and blue represent the samples in the rhizosphere of 2-y, 3-y, and 4-y transplanted ginseng seedlings, respectively.

PCoA ordination and Bray–Curtis distance matrix revealed that fungal communities obviously differed in the soils of ginseng seedlings of different ages and developmental stages (Figs. 3, S1). In vegetative stage, the first principal component axis (33.94% contribution) of the fungal communities in soils of 4-y seedlings markedly differed from those of the fungal communities in soils of 2-y seedlings. Moreover, the second principal component axis (17.55% contribution) of the fungal communities in soils of 2-year-old seedlings considerably differed from those of the fungal communities in the soils of 3-y seedlings (Fig.3E). At the flowering and fruiting stages, the first principal components (25.71% and 38.96% contributions, respectively) in the fungal communities in soils of 2-y ginseng seedlings significantly differed from those of the fungal communities in the soils of 3-y and 4-y seedlings (Figs. 3F, 3G). During root growth, the first principal component (26.97% contribution) of fungal communities in soils of 3-y seedlings considerably differed from those of fungal communities in soils of 2-y and 4-y seedlings, and the second principal component (14.93% contribution) in fungal communities in soils of 2-y seedlings differed from those of fungal communities in soils of 4-y seedlings (Fig. 3H). According to LEfSe analysis, fungal composition differed in the soils of ginseng seedlings of different ages and developmental stages (Fig. S3).

### Changes in the relative abundance of bacterial taxa

The relative abundance of bacterial groups changed in soils of ginseng plants of different ages and developmental stages (Fig. 4). The relative abundance of Chthoniobacteraceae, Chthonomonadales, Chthonomonadetes, *Chthoniobacter, Granulicella*, and *Blastocatella* declined with plant age in vegetative stage (Fig. 4A). The relative abundance of Arthrobacter, Brevundimonas, Micrococcaceae, Rhodobiaceae, Intrasporangiaceae, and Micrococcales was significantly higher in soils of 3-y and 4-y seedlings than that in soils of 2-y seedlings. The relative abundance of *Luteolibacter*, Phyllobacteriaceae, *Acidovorax, Moheibacter*, and Cytophagaceae in soils of 3-y and 4-y seedlings evidently decreased, and the abundance of Elusimicrobia and Armatimonadetes was markedly increased in the flowering stage (Fig. 4B). The relative abundance of *Bacillus*, Enterobacteriales, Enterobacteriaceae, *Brevundimonas*, and Anaerolineae was significantly higher in soils of 3-y and 4-y seedlings than that in soils of 2-y seedlings; additionally, the abundance of *Luteibacter*, Clostridia, and Clostridiales declined in soils of 3-y and 4-y seedlings in fruiting stage (Fig. 4C). The relative abundance of *Paralcaligenes*,Sphingomonadaceae, Saccharibacteria, *Sphingomonas*, and Alcaligenaceae significantly decreased with plant age, whereas the relative abundance of *Pandoraea*,Chlamydiales, and Chlamydiae was obviously higher in soils of 3-y and 4-y seedlings than that in soils of 2-y seedlings in root growth stage (Fig. 4D).

**Figure 4.**
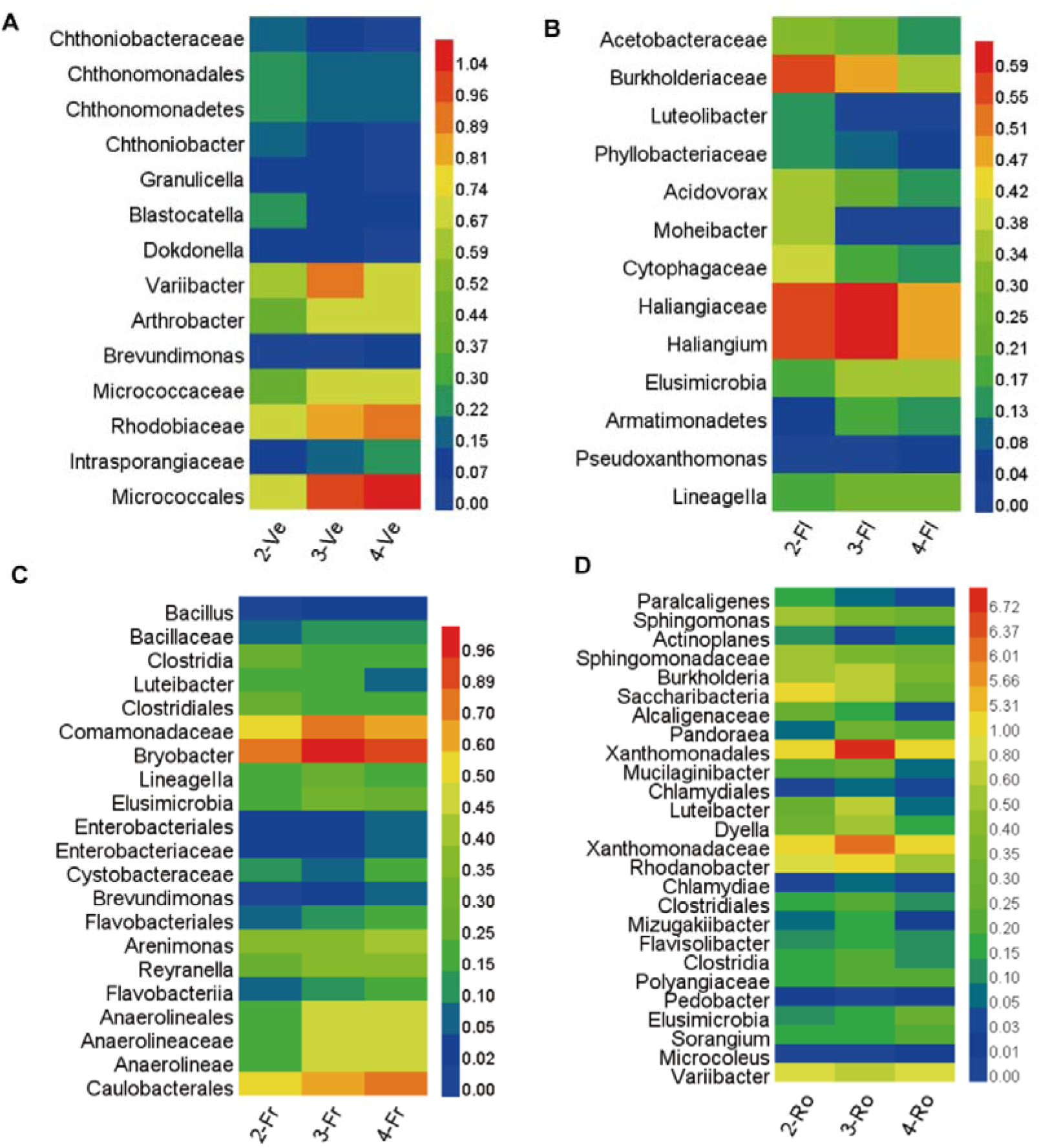
Relative abundance of the bacterial taxa detected by the linear discriminant analysis effect size (LEfSe) as biomarker. (A-D) represent the vegetative, flowering, fruiting, and root growth stages, respectively. 2-, 3- and 4- represent the samples in the rhizosphere of 2-y, 3-y, and 4-y transplanted ginseng seedlings. Data represent the mean values of *n* = 3.

### Changes in the relative abundance of fungal taxa

The relative abundance of fungal taxa changed in soils of ginseng plants of different ages and developmental stages (Fig. 5). The relative abundance of Cystofilobasidiales, Ophiostomataceae, *Ophiostoma*, Ophiostomatales, and Cystofilobasidiaceae significantly declined with seedling age, and the abundance of Pezizales, Cantharellales, *Dendryphion*, Pezizomycetes, and Tubeufiaceae was evidently higher in soils of 4-y seedlings than those in soils of 2-y and 3-y seedlings in the vegetative stage (Fig. 5A). The relative abundance of Microascales, *Helicoma*,and Tubeufiaceae was obviously higher in soils of 4-y seedling than those in soils of 2-y and 3-y seedlings in the flowering stage (Fig. 5B). The relative abundance of Tremellales, Acrospermales, *Occultifur, Acrospermum*, Cystobasidiales, Cystobasidiaceae, and Cystobasidiomycetes was markedly higher in the soils of 4-y seedlings than that in the soils of 2-y and 3-y seedlings in fruiting stage (Fig. 5C). The relative abundance of Zygomycota was obviously lower in soils of 3-y and 4-y seedlings than that in soils of 2-y seedlings, and the abundance of Tremellomycetes, Chytridiomycota, and Sordariales significantly increased in soils of 4-y seedlings than those in soils of 2-y and 3-y seedlings in root growth stage (Fig. 5D).

**Figure 5.**
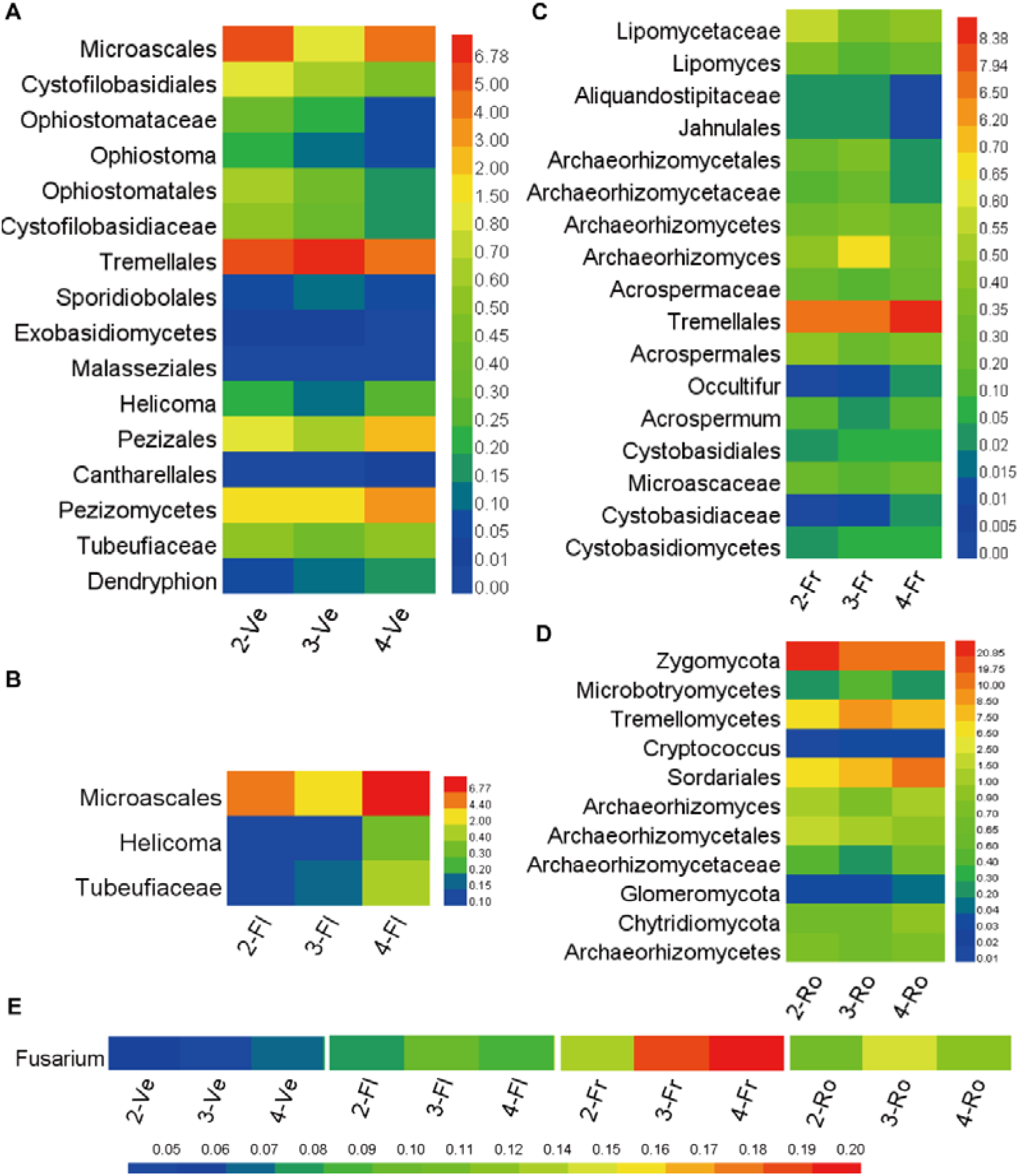
Relative abundance of the fungal taxa detected by LEfSe as biomarker and *Fusarium*. (A–D) represent the vegetative, flowering, fruiting, and root growth stages, respectively. (E) Relative abundance of *Fusarium*. Data are presented as mean ± SD (*n* = 3).

The relative abundance of *Fusarium* showed increasing trends in the soils of the adult ginseng plants under later developmental stages (Figs. 5E, S4). According to the high-throughput sequencing analysis, the relative abundance of *Fusarium* increased by 22.5%–25.0%, 35.7%–50.0%, and 18.2%–36.4% in soils of 3-y and 4-y seedlings in the flowering, fruiting, and root growth stages, respectively (Fig. 5E). The copy numbers of *Fusarium* also showed similar trends based on the analysis of quantitative PCR (Fig. S4). These results revealed that the relative abundance of pathogenic agent increased in soils during ginseng growth.

### *Bacillus subtilis* 50-1 were responsible for the biocontrol of *F. oxysporum*

Due to the relative abundance of pathogenic agent (*Fusarium*) increase in the ginseng soils, dual culture techniques were used to isolate microbial antagonists against *F. oxysporum* to control root rot. Antagonistic bacterium, namely, *B. subtilis* 50-1, was isolated and confirmed using a dual culture technique (Fig. 6). Strain 50-1 exhibited a broad spectrum of growth inhibition activity against *F. oxysporum*, thereby resulting in 67.8% inhibition percentage (Fig.6A). Strain 50-1 is a Gram-positive, oxidase- and catalase-positive, rod-shaped bacterial species (Fig.6B, Table S4). Analysis of the 16S rRNA sequences revealed that strain 50-1 belonged to *B*. *subtilis* with the bootstrap value of 100% (Fig. 6C). Strain 50-1 was further confirmed according to the genome sequencing. The complete genome was composed of a circular chromosome of 4,040,837 bp with 43.86% GC content (Fig. 6D, Table S5). The total numbers of genes were 4,193, which covered 88.6% of the genome encoding 3,176 proteins. The function of these annotated genes was mainly energy production and conversion, amino acid transport and metabolism, transcription, carbohydrate transport and metabolism, inorganic ion transport and metabolism, and signal transduction mechanisms based on the COG function classification (Fig. S5).

**Figure 6.**
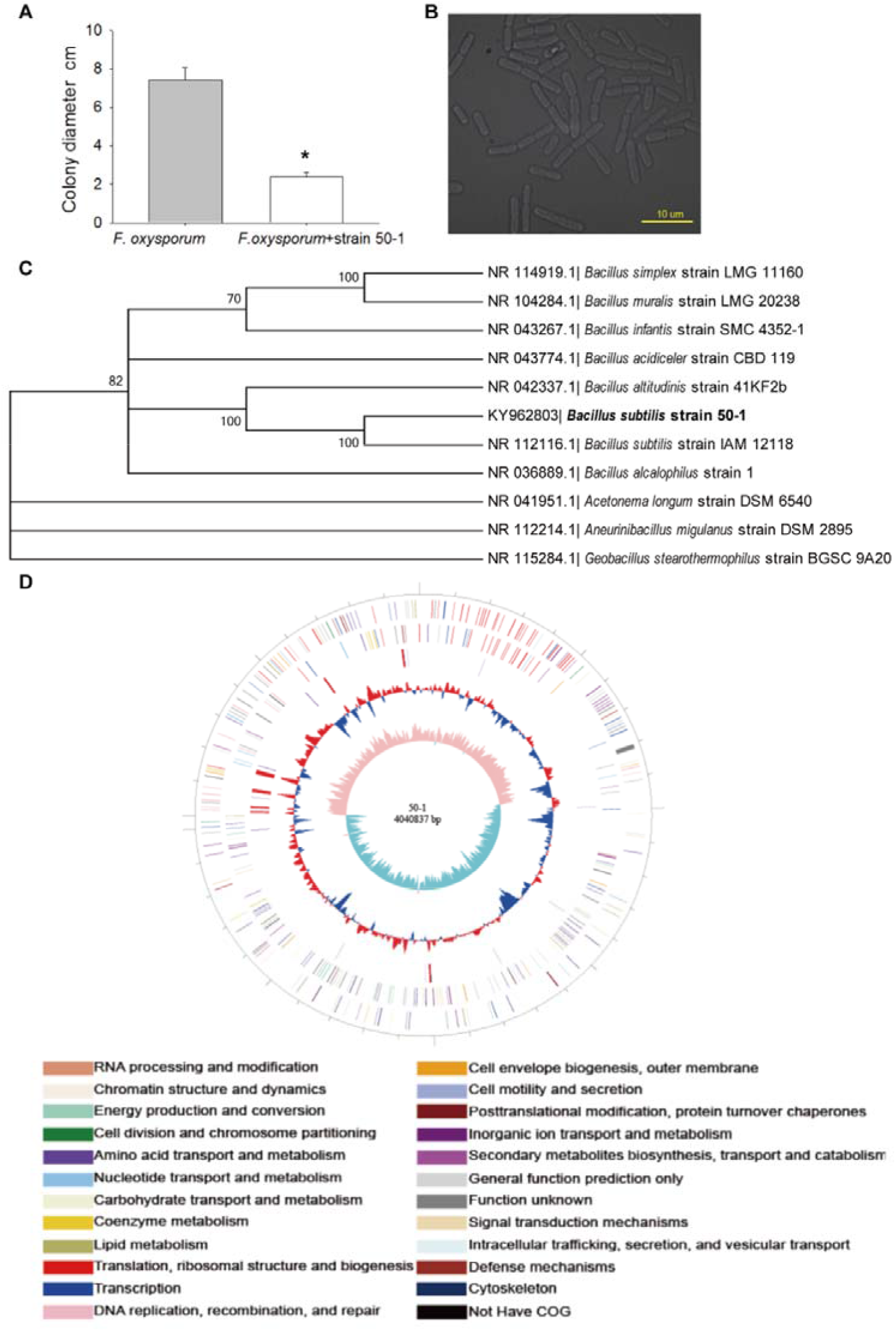
*Bacillus subtilis* 50-1 antagonized *Fusarium oxysporum*. (A) Colony diameter measured in a dual culture. (B) Morphological features of bacterium 50-1. (C) Relationships of 16S rRNA sequences between *B. subtilis* strain 50-1 (black body) and published 16S rDNA sequences. (D) Genome map of strain 50-1. The six circles (outer to inner) represent the scale line, forward strand CDSs (color by COG categories), reverse strand CDSs (color by COG categories), RNA genes, GC content, and GC skew. From outside to center: genome size, genes on the forward strand (color by COG categories), genes on the reverse strand (color by COG categories), RNA genes (tRNAs, orange; rRNAs, red), GC content (red and blue), and GC skew. Only bootstrap values higher than 70% are shown. Bars represent the mean ± SE (*n* = 3). Asterisks denote significant differences between the colony diameters of *F. oxysporum* and *F. oxysporum*+strain 50-1 at *P*< 0.05.

### Inoculation of biocontrol bacteria reduced ginseng death rate in a replanting soil

The pot experiment analysis revealed that ginseng death rate and the relative abundance of *Fusarium* significantly decreased by 63.3% and 46.1% in the replanting soils inoculated with strain 50-1, respectively (Fig. 7). Furthermore, the height and leaf area increased by 62.7% and 22.5%, respectively, after inoculation with strain 50-1 (Fig. S6). These results revealed that inoculation of biocontrol bacteria alleviated the ginseng replanting problem.

**Figure 7.**
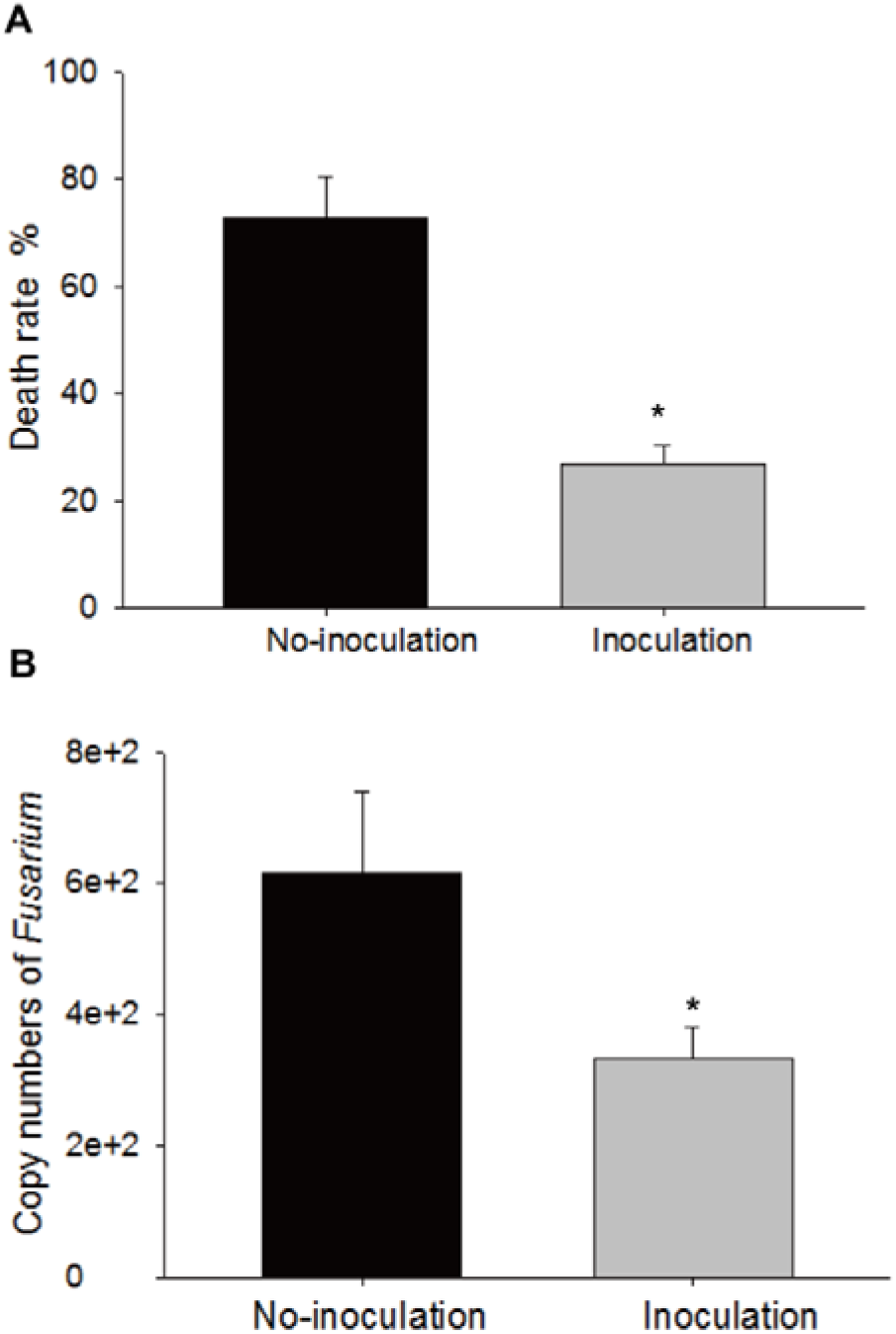
Inoculation of strain 50-1 in ginseng replanting soils. (A) Death rate of ginseng after inoculation with strain 50-1. (B) Copy numbers of *Fusarium* in soils. No-inoculation and inoculation represent the samples inoculated with cultures containing inactivated and activated strain 50-1, respectively. Data are presented as mean ± SD (*n* = 3). Asterisks denote significant di erences between the No-inoculation and inoculation at *P*< 0.05.

## Discussion

Replanting problem is a common and serious issue in cultivation of medicinal plants. Following replantation, *Rehmannia glutinosa* suffers from disease, and its biomass and tuber quality decline (Qi *et al*., 2009; Wu *et al*., 2011; Yang *et al*., 2011). The survival rate of ginseng seedlings is lower than 25% after replantation for three years (Zhao *et al*., 2001). Multiple factors have caused this replanting problem, and researchers suggested that changes in microbial communities influence soil health and crop yield (Bisseling *et al*., 2009; Bulgarelli *et al*., 2013). In addition, rhizospheric microbial community is governed by plant species and growth (Incroğlu *et al*., 2011). Thus, we hypothesized that changes in the rhizospheric microbial community were driven by ginseng seedlings of different ages and development stages, which caused the replanting problem of *Panax* plants. Reducing the occurrence of the disease is a useful strategy to alleviate the replanting issues. Biological control is one of the most remarkable potential approaches for plant disease control because of its safety and environmental friendliness.

Diversity in bacterial and fungal communities changed in the rhizosphere of ginseng seedlings of different ages and developmental stages. *H’* and Chao1 values revealed that bacterial diversity was obviously low, whereas fungal diversity was significantly higher in the rhizosphere of adult ginseng seedlings in the root growth stage. A similar study reported that increase in ginseng cultivation ages reduced bacterial diversity and increased fungal diversity (Xiao *et al*., 2016). Moreover, diversity of microbial community in the rhizosphere of *Pseudostellaria heterophylla* decreases with increased number of cropping years (Zhao *et al*., 2016). Additionally, the developmental stage of crops is an important driver of microbial community structure (Houlden *et al*., 2008). Plant ages and developmental stages alter the microbial diversity during growth. The diversity of soil microorganisms is critical in the maintenance of soil health and quality, and it serves as a sensitive bioindicator (He *et al*., 2008). The death rate of notoginseng and fungal diversity are significantly negatively correlated, which suggest that fungal diversity is a potential bioindicator of soil health (Dong *et al*., 2016). A relationship exists between microbial diversity and root disease suppression (Nitta, 1991; Workneh & van Bruggen, 1994). Mazzola (2004) reported that reduced soil microbial diversity is responsible for the development of soil-borne diseases. Reduced bacterial diversity in response to adult plants is possibly the key indicator of ecological variations and functional impairment in a root growth stage.

The bacterial and fungal compositions evidently differed in the rhizospheres of ginseng plants of different ages and developmental stages. Variation in bacterial and fungal community composition is observed during continuous cropping of notoginseng (Dong *et al*., 2016). Dynamics of microbial species is occurred in the rhizosphere during ginseng growth (Li *et al*., 2012). Sugiyama *et al*. (2014) reported that soybeans affected the bacterial communities under vegetative stage, with further alterations possibly occurring during later growth stages. The most divergent community structures occur in the young plant stage of tomato, and the flowering and senescence stages show increasingly similar community structure (Inceoğlu *et al*., 2011). The composition of microbial community was affected by plant age and development stage. In our study, the agents of beneficial microbes were relatively few, whereas those of harmful microorganisms increased with plant age. The agents of beneficial microbes include the microorganisms that degrade compounds (*Luteolibacter*, Cytophagaceae, *Luteibacter*, and *Sphingomonas*), and resist disease and improve growth (Sphingomonadaceae and Zygomycota). The agents of harmful microorganisms mainly infect plants (*Brevundimonas*, Enterobacteriaceae, *Pandoraea*, Cantharellales, *Dendryphion*, and Chytridiomycota). Another study has showed similar results, i.e., the population of beneficial microbe decreases, whereas that of harmful microorganisms increases with increasing number of cropping years (Zhao *et al*., 2016). Soil microbial community is an important bioindicator of soil function (Zuppinger-Dingley *et al*., 2014). Changes in functional groups revealed that micro-ecological environment was degraded in the rhizosphere with increasing age of ginseng.

Rhizospheric microbial community can be influenced by soil characteristics and plant species (Lauber *et al*., 2008; Berg and Smalla, 2009). In our study, the pH and available K and organic matter contents exhibited no significant difference in ginseng plants of different ages and developmental stages (Table S2). Plant species could be one of the most important factors that influence rhizobacterial communities (Micallef *et al*., 2009). Root exudates offer substrates for microbial metabolism and act as intermediates for biogeochemical reactions in rhizosphere (Uren, 2000; Berg and Smalla, 2009). In the early developmental stages of *Arabidopsis*, cumulative secretion levels of sugars and sugar alcohols are high, which enhanced the richness of rhizospheric microbial community (Chaparro *et al*., 2014). By contrast, root exudates contain allelochemical that disturbs the balance in a microbial community (Wu *et al*., 2015). These results imply that root exudates are one of the main drivers of changes in rhizospheric microbial communities during ginseng growth. Moreover, our results showed that the diversity of microbial communities markedly changed in the rhizosphere of ginseng plants with different ages in root growth stage. This phenomenon is possibly a result of different root types and root exudates. The composition of *Arabidopsis* root exudates changes throughout developmental gradient in plants, and tomato root exudates in reproductive stage are more phytotoxic than those in vegetative stage (Yu and Matsui, 1994; Chaparro *et al*., 2013). Ginseng root rapidly grows after cropping for three years before harvest, especially in root growth stage. Ginseng root growth stage is possibly characterized by a specific but different root exudation pattern that drives different bacterial communities.

Biological control using microbes is an environmentally friendly approach of controlling disease (Bargabus *et al*., 2003; Tjamos *et al*., 2005). In our study, we isolated *B. subtilis* 50-1 as an effective antagonist against *F. oxysporum*; the results of antagonist inoculation revealed that biocontrol bacterium reduced the ginseng morbidity and alleviated the replanting problem. *Bucillus* strains, as potent biological control agents of plant diseases, have been reported in many studies (Zouari *et al*., 2016; Shahzad *et al*., 2017). The biocontrol efficacy of *B. amyloliquefaciens* 54 against bacterial fruit blotch is evident in a greenhouse (Jiang *et al*., 2015). *B. megaterium* (B5), *B. cereus* sensu lato (B25), and *Bacillus* sp. (B35) display the highest antagonistic activity against *Fusarium verticillioides*, and antagonistic activity reveals that these strains produce glucanases, proteases, chitinases, siderophores, and auxins (Figueroa-López *et al*., 2016). Furthermore, data suggest that microbial strains producing bioactive constituents can help the inoculated plant to reduce the negative effects of pathogenesis and abiotic stresses (Shahzad *et al*., 2017). The microbial antagonists could be used for efficient control of plant pathogens and comprise an environmentally friendly approach. Additionally, genome sequence analysis provides insights into the pathways of functional bacteria and facilitates their exploration. The genome sequence analysis of *Ganoderma lucidum* revealed key genes encoding cytochrome P450s in secondary metabolism (Chen *et al.*, 2012). In our study, the genome sequence of strain 50-1 helps accelerate the development and application of biological inoculant.

## Supplementary data

The details were online.

**Table S1** Description of developmental stages of ginseng and the sampling times.

**Table S2** Soil chemical characteristics in soils of ginseng cropping.

**Table S3** Barcodes, numbers of bacterial and fungal sequences and average length in each sample.

**Table S4** Physiological and biological characteristics of Bacillus subtilis 50-1.

**Table S5** Genome features of *Bacillus subtilis* strain 50-1.

**Figure S1** Changes in bacterial and fungal community in rhizosphere of ginseng seedlings of different ages and developmental stages.

**Figure S2** LEfSe identified the taxa in the soil bacterial communities of plants of different ages in vegetative (A), flowering (B), fruiting (C), and root growth stages (D).

**Figure S3** LEfSe identified the taxa in the fungal communities of plants of different ages in vegetative (A), flowering (B), fruiting (C), and root growth stages (D).

**Figure S4** Relative abundance of *Fusarium* in the soils based on the analysis of real-time PCR.

**Figure S5** COG function classification in the Bacillus subtilis strain 50-1.

**Figure S6** The height (A) and leaf area (B) of ginseng seedlings after inoculation of
strain 50-1 in a replanting soil.

## Acknowledgements

This study was supported by grants from the National Science Foundation of China (81603238 and 81403053).

